# Dynamic regulation of endogenous transcription factor hubs at single-molecule resolution

**DOI:** 10.64898/2026.01.25.701606

**Authors:** Shawn Yoshida, Yanghao Zhong, Akiko Kumagai, William G. Dunphy, Shasha Chong

## Abstract

Eukaryotic transcription factors (TFs) form local, high-concentration hubs at specific genomic loci through dynamic, multivalent protein-protein interactions mediated by their low-complexity domains. The hub formation behavior plays an essential role in TFs’ transcriptional activation activities. Characterizing the dimensions, dynamics, and regulation of TF hubs requires high-resolution imaging of TFs in their native cellular environment, but much of such biophysical characterization remains missing. Here, we combined CRISPR/Cas9-mediated genome editing and advanced quantitative cell imaging, including single-molecule microscopy, to investigate the dynamic behaviors of the endogenous oncogenic fusion TF EWS::FLI1 in Ewing sarcoma cells. We found that endogenous EWS::FLI1 forms dynamic, sub-diffraction-limit hubs with mechanisms of dissolution that prevent the hubs from achieving macroscopic liquid-liquid phase separation. Hub formation is a neomorphic behavior of EWS::FLI1 that is not directly conferred by its parental proteins, EWSR1 and FLI1. We found that during mitosis, EWS::FLI1 hubs dissolve, but EWS::FLI1 molecules continue to dynamically bind and unbind mitotic chromosomes, revealing a role of EWS::FLI1 in mitotic bookmarking. Nascent RNA destabilizes EWS::FLI1 hubs on chromatin, but it does not affect the dimensions of the hubs. Finally, we visualized endogenous EWS::FLI1 hubs upon treatment with various compounds that were previously indicated to affect EWS::FLI1 function. We found that LY2835219 and trabectedin significantly alter the nuclear distribution of endogenous EWS::FLI1, disrupting and mislocalizing EWS::FLI1 hubs, respectively. This finding highlights the therapeutic potential of both compounds for Ewing sarcoma. Together, our results reveal new insights into the assembly and regulation of endogenous EWS::FLI1 hubs at an unprecedented resolution. The methodology developed here will be useful for characterizing the functional hubs of many regular and pathological TFs in the future.

## Introduction

DNA-binding transcription factors (TFs) are crucial regulators of eukaryotic transcription, comprising DNA-binding domains and transactivation domains that interact with other TFs and components of the transcriptional machinery to activate transcription. The transactivation domains usually contain intrinsically disordered low-complexity domains (LCDs) that do not fold into stable three-dimensional structures, thus are not amenable to investigation through the conventional structure-function framework. Recent work based on live-cell single-molecule imaging has revealed that TF LCDs undergo dynamic, multivalent, and selective protein-protein interactions, which drive the formation of high local concentration TF hubs at target genes. Under certain conditions, such as when LCDs are overexpressed, the hubs can develop into phase separated condensates. The multivalent interactions and hub formation behaviors of a TF enable the recruitment of more TFs^1,2^, RNA polymerase II^3–7^, and transcriptional cofactors to target genes^3,4,8^, leading to transcriptional activation. Notably, endogenous target genes often require optimal levels of multivalent LCD-LCD interactions for transactivation; both low and excessive levels of multivalent interactions can repress transcription^9^. This “Goldilocks behavior” applies to a variety of hub-forming transcriptional regulators^10–12^. TF LCDs are also frequently mutated in diseases, including many types of cancers, and the pathological implications of aberrant TF hubs have been documented in an increasing number of reports^13–17^. We note that here we use the term “hub” to refer to local high-concentration TF clusters mediated by LCD-LCD interactions. Historically, similar clusters have been described using various other terms, most often “condensate”, which was initially linked to liquid-liquid phase separation (LLPS) and now more broadly defined as membrane-less compartments that selectively concentrate biomolecules, regardless of whether LLPS is involved^18^. We adopt this broader definition of condensates, preserve the original terminology used in the cited studies, and use “assembly” as an inclusive, mechanism-agnostic term to describe LCD-mediated biomolecular clusters.

Despite the critical role of TF hubs in transcriptional control and disease mechanisms, many questions about them remain unanswered due to limitations of widely used imaging and sample preparation methods. Specifically, most existing works have investigated TF hubs using conventional, diffraction-limited microscopy techniques, which are unable to precisely characterize the dynamics, spatial distribution, and dimensions of sub-diffraction objects, thereby limiting the ability to link the physical properties of small TF hubs to their functional contributions^19^. In addition, many studies imaged fluorescently labeled proteins overexpressed in cells. While valuable insights have been generated in such settings, a significant caveat is that multivalent interactions and hub formation behaviors are sensitively dependent on protein concentrations^20,21^. Studies that perform extensive characterization of endogenous TF hubs using high-resolution imaging techniques are highly desirable but remain very limited in number^2,22^.

To fill in this gap, here we use single-molecule imaging methods, including photoactivated localization microscopy (PALM)^23^ and single-particle tracking (SPT)^24,25^, to characterize the dimensions, dynamics, and regulation of endogenous TF hubs in the cell. We use EWS::FLI1, an oncogenic fusion TF that is the hallmark of Ewing sarcoma, as the model TF in this study. It is encoded by a fusion gene produced by the t(11;22)(q24;q12) chromosomal translocation that fuses the intrinsically disordered LCD of EWSR1, an RNA-binding protein, with the well-structured DNA-binding domain of FLI1, a TF that plays roles in development and homeostasis^26^. We and others have previously reported that endogenous EWS::FLI1 forms local, high-concentration hubs at GGAA microsatellites via a combination of EWS::FLI1-DNA binding and homotypic, multivalent interactions of EWS LCD^1,27,28^. Yet, the dimensions of endogenous EWS::FLI1 hubs across the cell nucleus remain unquantified. It is also unknown whether the hub formation behavior is a property of either parental protein of EWS::FLI1 (EWSR1 or FLI1) or a neomorphic property of the oncogenic fusion. Another unexplored question is how EWS::FLI1 hubs are regulated during mitosis, when some TFs are evicted from chromatin as it compacts, and others remain bound to specific genomic loci^29^. The behaviors of TF hubs during mitosis impact how the gene expression program is restored after cell division^30^. In addition, although RNA is known to play a role in nuclear organization^31^ and modulate phase separation of specific transcription regulators, including MED1 and ENL^32,33^, it remains unknown whether RNA contributes to regulating endogenous TF hubs. Also unexplored is whether any compounds can modulate endogenous pathological TF hubs, such as EWS::FLI1 hubs, despite emerging research identifying small molecules that target condensates involved in transcriptional dysregulation^34^. We address these open questions by imaging endogenous EWS::FLI1 hubs at single-molecule resolution in Ewing sarcoma cells. This work sheds light on both the mechanistic underpinnings of transcriptional control and the therapeutic targeting of pathological TFs.

## Results

### Endogenous EWS::FLI1 forms sub-diffraction-limit hubs that undergo dynamic turnover

To quantify the dimensions of endogenous EWS::FLI1 hubs, we performed PALM on our previously established knock-in Ewing sarcoma cell line A673, where we fused a HaloTag to the C-terminus of endogenous EWS::FLI1 via CRISPR/Cas9-mediated genome editing^1^. With this labeling, we visualized the dynamic behaviors of EWS::FLI1 in its native cellular context with single-molecule resolution. PALM, based on the single-molecule localization technique, offers a spatial resolution of ∼20 nm with our imaging setup, significantly higher than diffraction-limited microscopy. A well-documented issue with traditional PALM is spurious clustering caused by repeated appearances of the same molecule across multiple frames, due to factors including frame discretization of a single emission event, variable photobleaching times, and photoblinking^35–41^. Among different correction methods, pair-correlation PALM (PC-PALM) is known to faithfully report true clustering, making it an excellent tool for probing the presence of clusters and their length scale^40,42,43^. We applied PC-PALM to the raw single-molecule localizations of EWS::FLI1^44–46^ to determine whether the apparent clustering of localizations represents true EWS::FLI1 clusters or artifacts. We observed statistically significant clustering of EWS::FLI1 with an average cluster correlation length of *ξ* = 29.87 ± 1.89 *nm* (hereafter, mean ± SEM) (Figure 1a). The correlation length is known to provide an estimate of average cluster radius, though this is only an approximation and should not be interpreted as an exact measure^47^ (Methods). Our measurement suggests that endogenous EWS::FLI1 hubs have an average diameter of ∼60 nm (2*ξ*). In addition, while PC-PALM can robustly identify various molecule density fluctuations, including clustering, it cannot be used to generate a reconstructed image of the whole nucleus or to recover the distribution of cluster sizes across the nucleus. Nevertheless, PC-PALM confirms that endogenous EWS::FLI1 hubs are smaller than the diffraction limit of light and justifies the need for super-resolution microscopy for their characterization.

**Figure 1.**
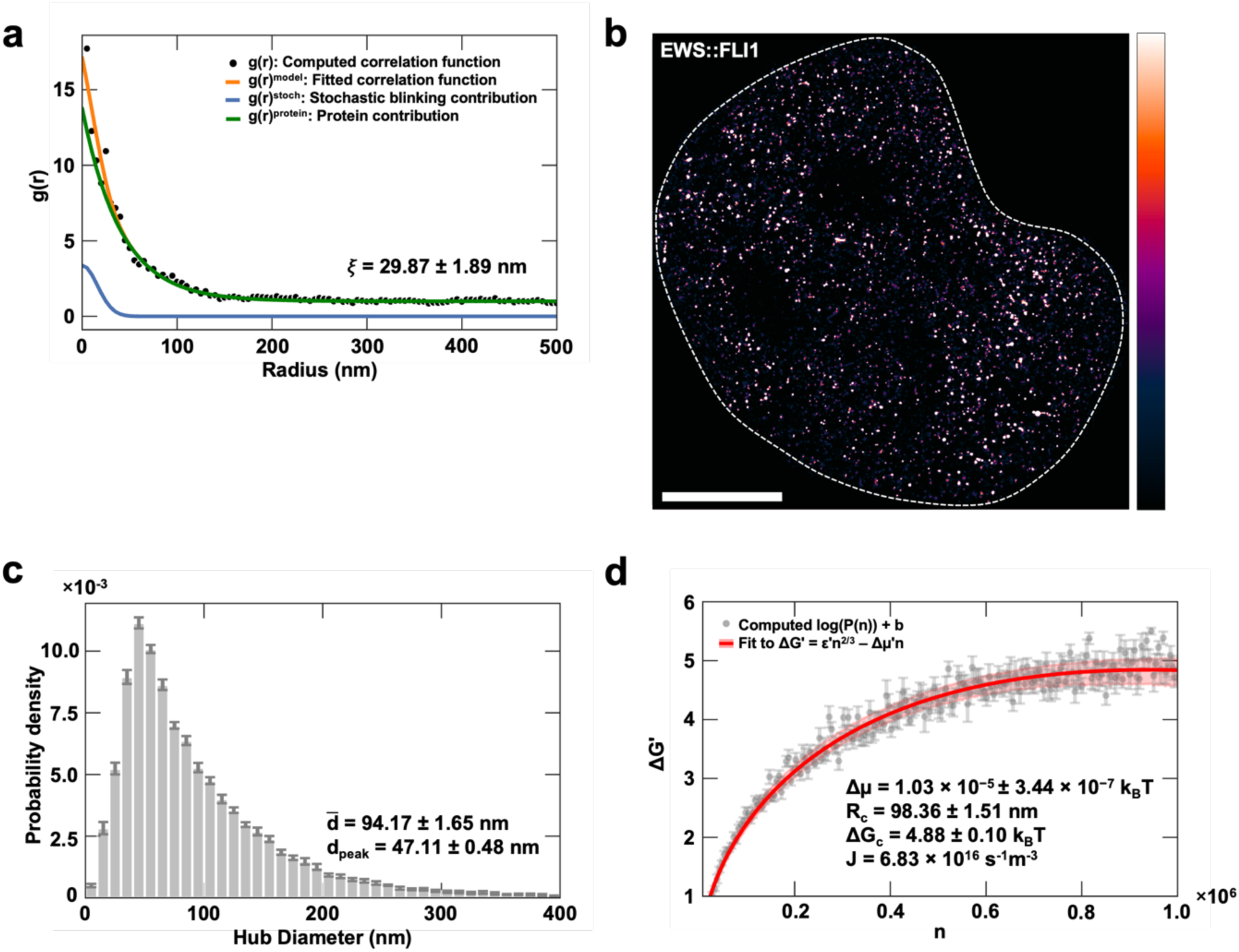
Super-resolution microscopy characterizes the dimensions of endogenous EWS::FLI1 hubs. (a) Pair-correlation analysis implemented as described previously^46,47^ characterizes the spatial distribution of EWS::FLI1 hubs. The computed pair-wise autocorrelation function of the raw PALM localizations (black dots) was fit to a model that accounts for the contributions of both true protein clustering and the multiple appearances of the same fluorophores due to stochastic blinking (orange curve). The protein correlation term and stochastic blinking correlation term are shown as green and blue curves, respectively. The correlation length of protein clusters, ξ, is proportional to the cluster size and 2ξ can be used as an estimate of average cluster diameter. Statistically significant clustering was observed with 2ξ = 59.74 ± 3.77 nm. (b) Photoblinking-corrected PALM image of a knock-in A673 cell expressing endogenous EWS::FLI1-Halo (labeled with the Halo ligand PA-JF549) reveals a punctate distribution of EWS::FLI1 in the nucleus, which forms numerous sub-diffraction-limit hubs. A white dashed contour outlines the cell nucleus. Scale bar is 5 µm. (c) The distribution of effective hub diameters generated by clustering the photoblinking-corrected localizations with DBSCAN^48^. (d) Plot of −log(*P*(*n*)) vs n fitted with *ɛ*^′^*n*^2/3^ − *Δμ*^′^*n* yields values of *ɛ*^′^ and *Δμ*^′^ used to calculate the free energy barrier to nucleation and critical cluster size. Data is collected from 10 cells. SEM represents the uncertainty.

To extract the true EWS::FLI1 localizations for image reconstruction, we applied statistical corrections for photoblinking to the raw single-molecule localizations. Briefly, the blinking behavior of PA-JF549 in A673 cells was characterized within the framework of a stochastic model, and the resulting rate parameters were used to filter out repeated localizations arising from blinking and frame-discretization artifacts from raw PALM datasets (Methods). The resulting PALM reconstruction shows the punctate distribution of endogenous EWS::FLI1 (Figure 1b). Density-based spatial clustering of applications with noise (DBSCAN) was applied to the corrected localizations to identify individual EWS::FLI1 hubs^49^. We then plotted the distribution of hub diameters and found that the hubs have a mean diameter of 47.11 ± 0.48 *nm* and an estimated mode diameter of 94.17 ± 1.65 *nm* (Figure 1c). This is consistent with the PC-PALM result, confirming that endogenous EWS::FLI1 forms sub-diffraction hubs.

We next extracted parameters governing the energetics and kinetics of endogenous EWS::FLI1 hub formation by analyzing the hub size distribution through the lens of classical nucleation theory, a powerful theoretical framework for predicting nucleation kinetics. While nucleation classically describes the first step in the formation of a new thermodynamic phase, its use here does not imply that EWS::FLI1 undergoes LLPS; rather, we apply it as a general mathematical framework for describing the stochastic formation of clusters as established in the field^50^. In the classical, mean-field treatment of homogeneous nucleation, Δ*G*, the free energy change for the nucleation of a cluster of size *n*, can be compactly written as^51^

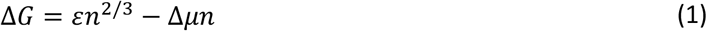

The first term in Eq. 1 represents the contribution from the surface energy to Δ*G*, i.e. the energetic cost of maintaining the interface between the cluster and the surrounding environment, encapsulated by the prefactor, *ɛ*. The energetic cost depends on factors including the molecule size, the geometry of the interface, and the magnitude of interfacial energy, and it always increases with the cluster size. The second term in Eq. 1 represents the contribution from the bulk free energy to Δ*G*, i.e. the change in free energy due to the difference in chemical potential, Δ*μ*, between the inside of the cluster and the surrounding environment, where Δ*μ* = *μ* − *μ*^∗^ = *k*_*B*_*T* log(*c*⁄*c*^∗^) . *c*⁄*c*^∗^ is the ratio of the actual in-cluster concentration to the equilibrium or saturation concentration. While the concept of a saturation concentration is often associated with LLPS, here it instead denotes the concentration above which the formation of stable clusters becomes favorable, independent of macroscopic phase separation. In subsaturated systems (Δ*μ* < 0), both the surface and bulk energy contributions increase with cluster size, making any clusters that spontaneously form due to thermal fluctuations tend to dissipate. Conversely, in supersaturated systems (Δ*μ* > 0), the surface energy still increases with increasing cluster size, but the bulk energy decreases with increasing cluster size. As a result, while some clusters still tend to dissipate, others can overcome the free energy barrier and continue to grow larger. *ɛ* and Δ*μ* together set the free energy barrier to nucleation, Δ*G*_*c*_ , and the corresponding critical radius, *R*_*c*_ , which represents the minimum radius above which it is thermodynamically favorable for a cluster to continue growing rather than being driven to dissipation. *ɛ*, Δ*μ*, *R*_*c*_, and Δ*G*_*c*_, can all be determined from the distribution of cluster sizes, *P*(*n*), with^52^

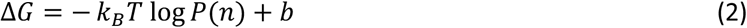

Here, *b* is a self-consistent term that ensures the normalization of *P*(*n*). The derivation of Eq. (1), along with expressions for *R*_*c*_and Δ*G*_*c*_, is included in the Methods.

We fitted Eq. (2) against the EWS::FLI1 hub size distribution (Figure 1d), and remarkably, the shape of the distribution matches what would be expected of a supersaturated system in the framework of the classical nucleation theory. The difference in chemical potential (Δ*μ*) is 1.03 × 10^−5^*k*_*B*_*T*, whose positive sign suggests that at endogenous expression levels, EWS::FLI1 is supersaturated. The critical radius (*R*_*c*_) is 98.36 ± 1.51 *nm*, and the free energy barrier to nucleation (Δ*G*_*c*_ ) is 4.88 ± 0.10 *k*_*B*_*T*. The relatively modest free energy barrier suggests that EWS::FLI1 hub nucleation is a frequent and reversible process.

Nucleation of a supersaturated state is typically transient and followed by cluster growth. As such, if we consider supersaturated nucleation of EWS::FLI1 hubs in isolation, an increasing number of hubs would overcome the free energy barrier and continue to grow beyond the critical radius, eventually becoming large LLPS droplets. However, the actual size distribution of EWS::FLI1 hubs is stable over time in the cell, suggesting that a balance is maintained between nucleation of new hubs and dissolution of hubs larger than the critical size. Using the Zeldovich form of the steady-state nucleation rate equation^53^ along with the energetic parameters extracted earlier, we estimate that EWS::FLI1 hubs have a nucleation rate of 6.83 × 10^16^ *m*^−3^*s*^−1^ (Methods). This rate should match the bulk dissolution rate to maintain the observed steady-state hub size distribution. As such, approximating the volume of an A673 nucleus as 500 *μm*^3^ from our imaging data, the dissolution rate is about 34 hubs per second. While the mechanism underlying hub dissolution remains unknown, this estimate sets the timescale for dissolution, allowing us to narrow down possible dissolution pathways. Previously reported processes of regulating intrinsically disordered region (IDR)-mediated assemblies or condensates, including chaperone-mediated dissolution^54–56^, phosphorylation-mediated dissolution^57–59^, and competition between multivalent interactions and DNA-protein binding^60^, can occur at rates near 34 hubs per second. In contrast, slower processes, including targeted proteasomal degradation, are unlikely to operate on this timescale.

### EWS::FLI1 hub formation is a neomorphic behavior not shared by EWSR1 or FLI1

Many fusion oncoproteins resulting from chromosomal translocations can phase separate or form biomolecular assemblies via multivalent interactions^61–63^. EWS::FLI1 is one of such oncoproteins. It remains unexplored if the assembly formation ability of a fusion oncoprotein is directly conferred by any of the parental proteins or a neomorphic property of the fusion. To address this question for EWS::FLI1, we used PALM, along with the above-described photoblinking corrections, to visualize the spatial distributions of EWSR1 and FLI1 in the cell and compare them with the distribution of EWS::FLI1. It was crucial to image both parental proteins at native expression levels, given that multivalent interactions and hub formation behaviors, if present, are highly dependent on protein concentration. To fluorescently label EWSR1, we engineered the A673 line with CRISPR/Cas9-mediated genome editing to fuse a HaloTag to the N-terminus of endogenously expressed EWSR1. Halo-tagging endogenous EWSR1 did not affect its expression levels (Figure 2a). Because FLI1 is not expressed in A673, we could not image endogenous Halo-tagged FLI1 in this cell line^64^. The A673 line is also not ideal for studying exogenously expressed FLI1, whose interaction behaviors are likely affected by the presence of endogenous EWS::FLI1 that may compete with FLI1 for the same binding sites on chromatin^65^. We instead opted to express exogenous Halo-tagged FLI1 in U2OS cells with no endogenous EWS::FLI1 or FLI1^66,67^. To ensure that we measure the intracellular distribution of FLI1-Halo at expression levels comparable to those of endogenous EWS::FLI1-Halo in A673 cells, we performed PALM for FLI1-Halo in U2OS and EWS::FLI1-Halo in A673 cells until complete depletion of fluorescence, i.e., no more molecules could be detected, quantified the total number of single-molecule localizations detected throughout the PALM movie as a measure of the protein’s expression level, and selected only the U2OS cells expressing FLI1-Halo at levels similar to endogenous EWS::FLI1-Halo in A673 cells for downstream analysis of FLI1 distribution (Figure 2b).

**Figure 2.**
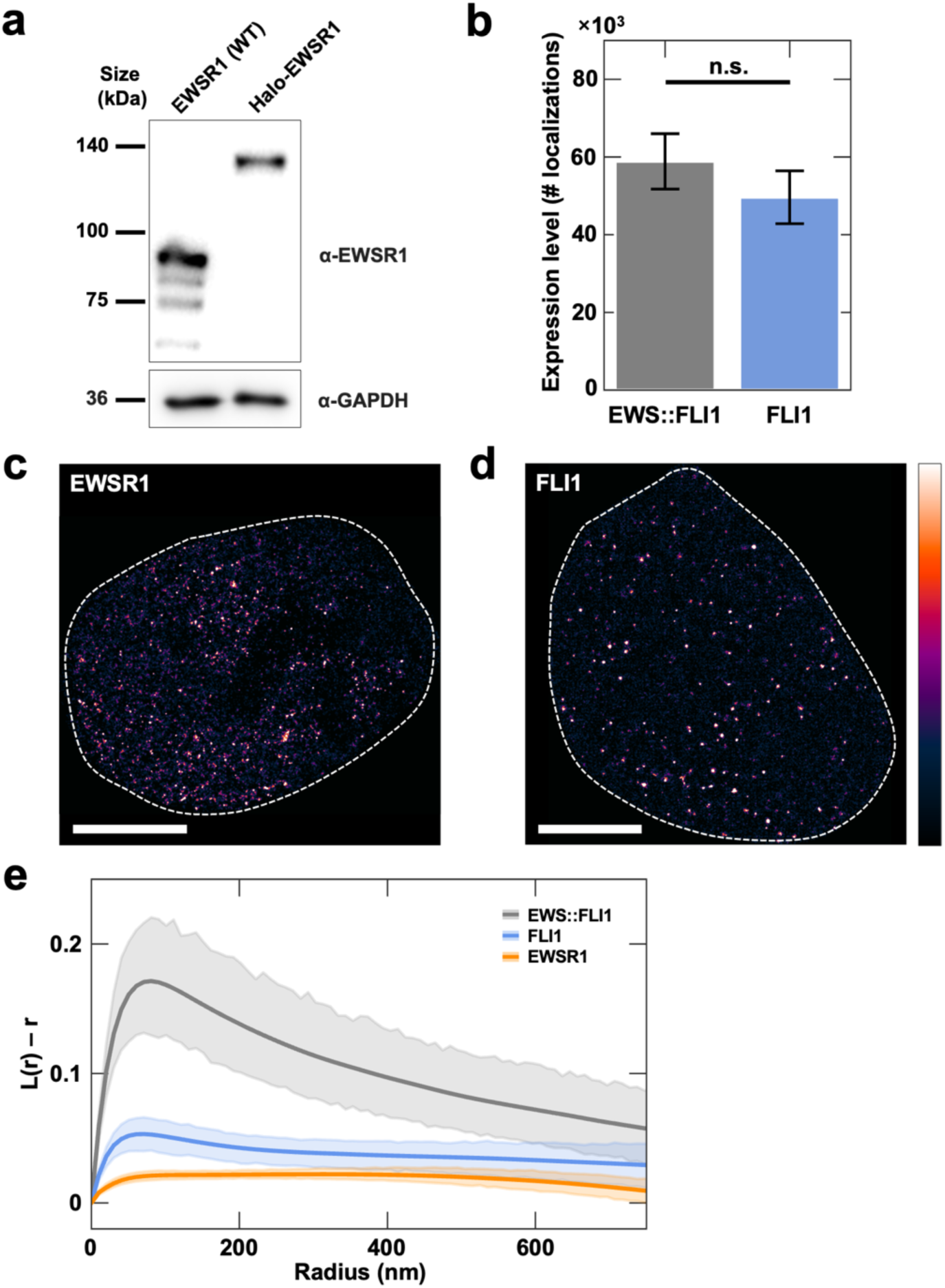
EWS::FLI1 has a stronger hub-forming ability than its parental proteins. (a) Western blot of EWSR1 and GAPDH from wild-type (WT) A673 and Halo-EWSR1 knock-in A673 cell lines. (b) Plot showing no statistically significant difference in the expression levels of endogenous EWS::FLI1-Halo in knock-in A673 cells and exogenous FLI1-Halo in U2OS cells (*p* = 3.89 × 10^−1^ , Student’s t-test). The expression level is quantified by the total number of single-molecule localizations detected throughout the PALM movie of each cell. 10 cells were analyzed for each condition. Error bars represent SEM. Not statistically significant is abbreviated as n.s. (c-d) Photoblinking-corrected PALM images of a knock-in A673 cell expressing endogenous Halo-EWSR1 (c) and a U2OS cell transiently expressing FLI1-Halo at a level comparable to endogenous EWS::FLI1-Halo in A673 (d). The cells were stained with the Halo ligand PA-JF549. Both EWSR1 and FLI1 are less clustered than EWS::FLI1, though they still form puncta. A white dashed contour outlines the cell nucleus. Scale bars are 5 µm. (e) Ripley’s L(r) -r curves show that over relevant length scales, EWS::FLI1 is more clustered than both EWSR1 and FLI1. 10 cells were analyzed for each condition. The semi-transparent regions represent 95% confidence intervals.

PALM revealed that both EWSR1 and FLI1 have punctate distributions in the cell nucleus when expressed at endogenous levels (Figure 2c and 2d). However, their puncta are significantly less pronounced than the hubs formed by endogenous EWS::FLI1. We quantified the protein distribution by computing Ripley’s L function, *L*(*r*), a spatial descriptive statistic that determines whether a set of points has a random, dispersed, or clustered distribution at a given length scale^68^. Specifically, we compared plots of *L*(*r*) − *r* vs *r* for the localizations of EWS::FLI1, EWSR1, and FLI1 (Figure 2e), where randomly distributed localizations would yield a *L*(*r*) − *r* value of 0, and the more punctate the localization distribution, the larger the magnitude of *L*(*r*) − *r*. As shown in Figure 2e, EWS::FLI1 has a significantly more punctate distribution than both its parental proteins, EWSR1 and FLI1. This result suggests that the hub-forming ability of EWS::FLI1 is neomorphic instead of being directly conferred by either of its parental proteins alone. Although here we focus on a specific oncogenic fusion TF, EWS::FLI1, it is possible that some other aberrant fusion proteins resulting from chromosomal translocation similarly have their intracellular distributions altered from their parental proteins and form neomorphic assemblies, and the assembly formation behaviors play a role in the oncogenesis of different cancer types beyond Ewing sarcoma.

### EWS::FLI1 hubs dissolve during mitosis, and EWS::FLI1 dynamically interacts with mitotic chromatin

Since IDR-mediated biomolecular assemblies are involved in many cellular processes, their evolution during mitosis can impact many cellular functions. Despite this, the mitotic behaviors of only a handful of assembly-forming proteins have been investigated thus far; for example, mitotic hyperphosphorylation of the nucleolar proteins Ki-67 and NPM1 is reported to alter the charge blockiness of their IDRs, i.e. whether charged amino acids are arranged in large blocks or randomly dispersed, thereby modulating their condensation levels^69^. However, mitotic regulation of TF hubs remains unexplored. To fill this gap, we tracked the fate of endogenous EWS::FLI1 hubs throughout the cell cycle and compared the interaction and diffusion dynamics of EWS::FLI1 during mitosis and interphase. Since the association of many TFs with mitotic chromatin is known to be affected by cell fixation^70^, here we imaged EWS::FLI1 hubs in live knock-in A673 cells that express endogenous EWS::FLI1-Halo. We treated the cells with the anti-mitotic agent nocodazole to synchronize them in the cell cycle. Meanwhile, we stained the cells with a DNA dye (Hoechst 33342) that reveals the morphology of chromatin or chromosomes, enabling identification of specific phases of the cell cycle. Using live-cell confocal fluorescence microscopy, we found that endogenous EWS::FLI1 hubs dissolve upon mitotic entry, with no visible hubs detected during prometaphase, metaphase, or anaphase. However, during telophase, the hubs began to reform, coinciding with chromatin decondensation (Figure 3a). Interestingly, despite the absence of visible EWS::FLI1 hubs through most of mitosis, EWS::FLI1 remained weakly enriched on mitotic chromatin, consistent with previous report by Teves et al. that many TFs bind to mitotic chromatin and serve as “mitotic bookmarks”, which maintain established transcriptional programs throughout the cell cycle and restart them in daughter cells after mitosis^70^. Mitotic bookmarking by EWS::FLI1 is likely especially important in Ewing sarcoma cells, given its role in maintaining the oncogenic transformation phenotype.

**Figure 3.**
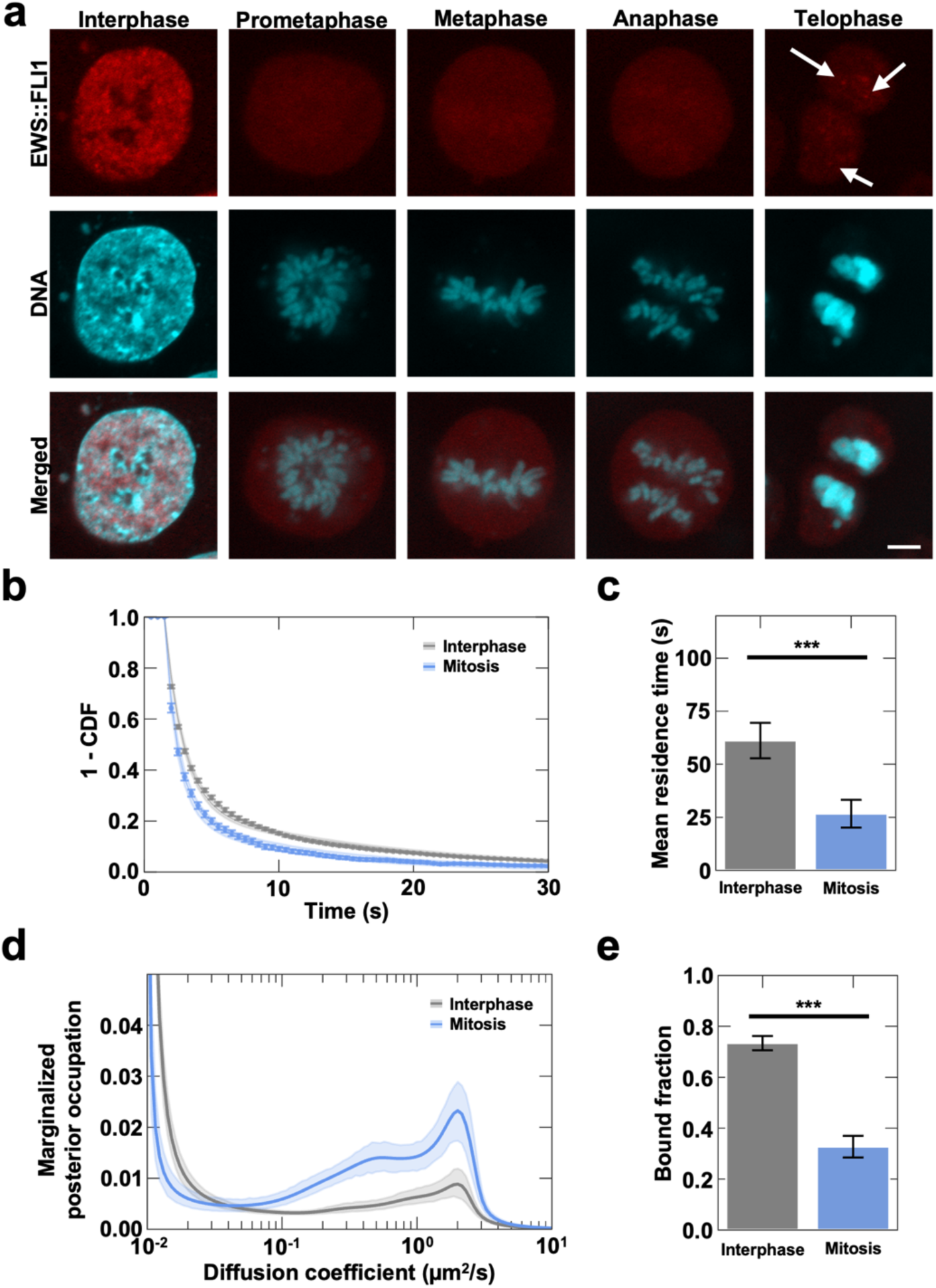
Interaction dynamics of EWS::FLI1 during mitosis. (a) Live-cell confocal images of the mitotic localization of endogenous EWS::FLI1-Halo in the knock-in A673 cell line. The cells were labeled with the Halo ligand JFX646 and the DNA stain Hoechst 33342. EWS::FLI1 hubs dissolve during mitosis and reform during telophase. Arrows highlight the reformed hubs. Scale bar is 5 µm. (b) Survival curves (scatter plots) were generated based on the binding residence times of individual EWS::FLI1 molecules on chromatin during interphase and mitosis, which were fit as described in the Methods (line plots). (c) The plot of the mean residence times shows that EWS::FLI1 binds more stably during interphase than during mitosis (***, *p* = 1.46 × 10^−4^, Mann-Whitney U test). For each condition, 21 cells were measured in three independent imaging sessions, each session performed on a different day. (d) EWS::FLI1 diffusion profiles show comparable diffusion coefficients during interphase and mitosis. Diffusion coefficients were extracted as the local maxima of the respective diffusion coefficient distributions. The semi-transparent regions represent 95% confidence intervals. (e) Plot of bound fractions shows that a greater fraction of EWS::FLI1 is bound during interphase than during mitosis (***, *p* = 2.70 × 10^−10^, Welch’s t-test). EWS::FLI1 molecules were considered bound if they had a diffusion coefficient smaller than the threshold value 0.1μm^2^/*s*. For each condition in (d) and (e), 30 cells were measured in three independent imaging sessions, each session performed on a different day. Error bars represent SEM in (c) and (e).

We next used SPT to compare the chromatin binding dynamics of endogenous EWS::FLI1 during interphase and mitosis in the EWS::FLI1-Halo knock-in A673 cells^1^. Our SPT protocol has been described in detail previously^71^. Briefly, we performed single-molecule imaging with a relatively long exposure time (500 ms, “slow” SPT) such that quickly diffusing molecules are blurred out and bound molecules are clearly resolvable. We then fitted the survival curve generated using the binding residence times of all the detected molecules in a cell to a two-component model of dissociation that accounts for both specific and nonspecific binding (Figure 3b)^71^. This allowed us to extract the mean residence times of EWS::FLI1 specifically bound to chromatin. We found that EWS::FLI1 binds to chromatin with a mean residence time of 61.15 ± 8.36 *s* during interphase and 26.71 ± 6.55 *s* during mitosis (Figure 3c). The fact that EWS::FLI1 binds to mitotic chromatin more transiently coincides with the dissolution of EWS::FLI1 hubs during mitosis. This coincident timing is consistent with our previous finding that hub formation of EWS::FLI1 during interphase stabilizes its chromatin binding through multivalent LCD-LCD interactions in the hubs^1^, further supporting the role of the multivalent interactions in regulating TF-chromatin binding.

We also compared the diffusion dynamics of endogenous EWS::FLI1 in the EWS::FLI1-Halo knock-in A673 cells during interphase and mitosis using SPT. To ensure the tracking of both bound and freely diffusing molecules in single-molecule images, we used a much shorter exposure time (5 ms, “fast” SPT) combined with stroboscopic illumination^25,72^. We then analyzed the resulting single-molecule trajectories using the state-array SPT (saSPT) algorithm, which extracts the diffusion coefficient distributions of a target protein without *a priori* knowledge of the number or identity of specific diffusing states of the protein^73^ (Figure 3d). We found that while the characteristic diffusion coefficient of freely diffusing EWS::FLI1 varies minimally with the cell cycle, with its value being 1.96 ± 0.02 *μm*^2^/*s* during interphase and 2.00 ± 0.04 *μm*^2^/*s* during mitosis, the bound fraction of EWS::FLI1 significantly varies with the cell cycle, i.e., 73.41 ± 2.80% during interphase and 32.73 ± 4.25% during mitosis (Figure 3e). The decreased bound fraction during mitosis coincided with the dissolution of EWS::FLI1 hubs, the formation of which during interphase likely helps to retain EWS::FLI1 on chromatin.

Together, these results suggest that EWS::FLI1 hub assembly and disassembly are regulated throughout the cell cycle. The dissolution of hubs during mitosis coincides with decreased binding stability and bound fraction of EWS::FLI1 to chromatin. Nonetheless, during mitosis, a fraction of EWS::FLI1 remains bound to chromatin, suggesting it may function as a mitotic bookmark^74,75^.

### RNA modulates the hub assembling stability of EWS::FLI1 but not its spatial distribution

It has been increasingly appreciated that RNA can play a role in regulating biomolecular assemblies. For example, Henninger et al. recently proposed an RNA-mediated feedback mechanism for Mediator condensates, where condensate formation is promoted by low levels of RNA when genes start to be transcribed, and the condensates are dissolved upon accumulation of the transcribed RNA^33^. However, it remains unknown whether this model is applicable to endogenous TF hubs that often form at actively transcribed genes. To tackle this question, we investigated the role of RNA in the organization and dynamics of EWS::FLI1 hubs in the knock-in A673 cells that express endogenous EWS::FLI1-Halo^1^ by imaging the hubs before and after we treated the cells with triptolide, a transcription initiation inhibitor that induces the proteasomal degradation of Rpb1, the largest subunit of RNA polymerase II^76^. Triptolide is known to globally inhibit transcription within minutes, but additional time is needed for all nascent RNAs to be degraded or diffuse away from their gene loci after transcriptional inhibition. To determine the time needed to deplete nascent RNA from the target genes of EWS::FLI1, we performed intron RNA fluorescence in situ hybridization (FISH)^70^ to detect nascent RNA of two well-characterized target genes of EWS::FLI1, *ABHD6* and *CAV1*^77^, at multiple timepoints after we treated the cells with 1 µM triptolide dissolved in DMSO. We found that the nascent RNA of both genes became undetectable two hours after triptolide treatment (Figure 4a). We then performed PALM to visualize the distribution of endogenous EWS::FLI1 after treating the cells with triptolide for two hours (Figure 4b). The *L*(*r*) − *r* vs *r* plots showed that the spatial distribution of EWS::FLI1 in the nucleus after the triptolide treatment does not differ significantly from after the treatment of DMSO only for two hours (Figure 4c). By measuring the EWS::FLI1 hub dimensions after the triptolide treatment, we found that hubs have a mean diameter of 46.54 ± 0.42 *nm* and an estimated mode diameter of 90.03 ± 2.25 *nm* (Figure S1a), which are not significantly different from those in cells treated with DMSO only (Figure 1c). These results suggest that RNA does not play a significant role in maintaining the assembly of EWS::FLI1 hubs.

**Figure 4.**
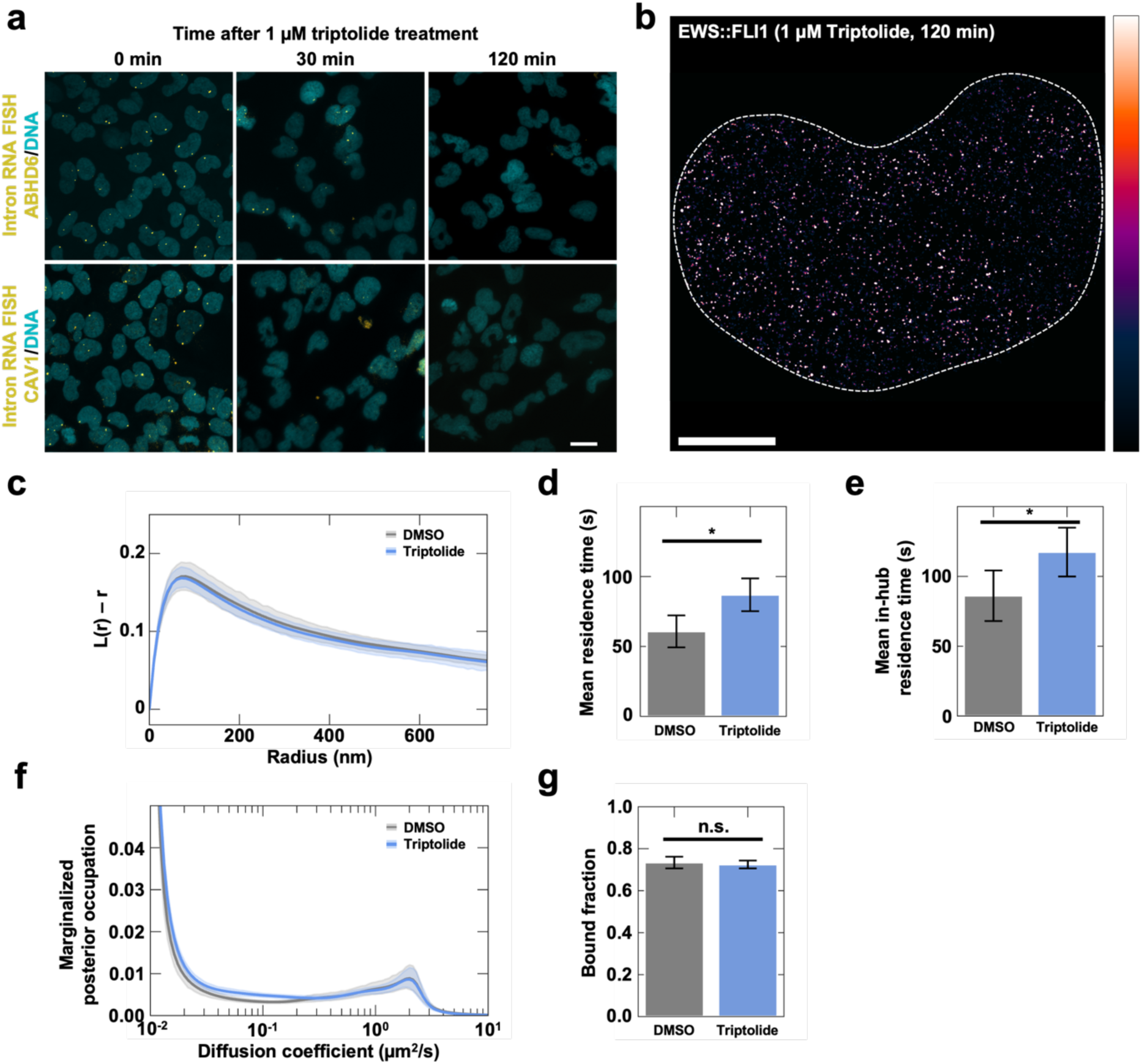
Transcriptional inhibition stabilizes EWS::FLI1 binding to hubs. (a) Confocal microscopy time-lapse of intron RNA FISH targeting *ABHD6* transcripts (FISH probes labeled with Quasar 670, yellow) or *CAV1* transcripts (FISH probes labeled with Quasar 570, yellow), and DNA (labeled with Hoechst 33342, cyan) in the knock-in A673 cells expressing endogenous EWS::FLI1-Halo after the treatment with 1 µM of triptolide. Scale bar is 20 µm. (b) Photoblinking-corrected PALM image of a knock-in A673 cell expressing endogenous EWS::FLI1-Halo (labeled with the Halo ligand PA-JF549) after transcriptional inhibition with triptolide. Scale bar is 5 µm. (c) Ripley’s L(r) – r curves show that the punctate distribution of EWS::FLI1 does not differ significantly between the DMSO-treated and triptolide-treated cells. (d-e) Plots of the mean residence times of EWS::FLI1 show that it binds more stably across the chromatin (*, *p* = 2.06 × 10^−2^, Mann-Whitney U test) (d) and within its hubs (*, *p* = 2.21 × 10^−2^, Mann-Whitney U test) (e) upon transcriptional inhibition. For each condition, 21 cells were measured in three independent imaging sessions, each session performed on a different day. (f) EWS::FLI1 diffusion profiles show comparable diffusion coefficients between the DMSO-treated and triptolide-treated cells. (g) Plot of bound fractions shows comparable bound fractions in DMSO-treated and triptolide-treated cells (n.s., *p* = 1.96 × 10^−1^ , Mann-Whitney U test). For each condition in (f) and (g), 30 cells were measured in three independent imaging sessions, each session performed on a different day. The semi-transparent regions represent 95% confidence intervals in (c) and (f). Error bars represent SEM in (d), (e), and (g).

We next asked whether the dynamics of EWS::FLI1 hub assembly are dependent on the presence of RNA. To this end, we measured the mean chromatin-binding residence times of EWS::FLI1 after triptolide or DMSO-only treatment using slow SPT. We found that in cells treated with DMSO only, EWS::FLI1 binds to chromatin with a mean residence time of 60.77 ± 11.44 *s*. The residence time increases to 86.93 ± 11.71 *s* in the cells treated with triptolide (Figure 4d). Similarly, the mean residence time of EWS::FLI1 in its hubs increases from 86.09 ± 18.13 *s* to 117.37 ± 17.48 *s* upon triptolide treatment (Figure 4e)^1^. This result suggests that the presence of RNA in EWS::FLI1 hubs facilitates dissociation of EWS::FLI1 from the hubs, i.e., increases its dissociation rate constant *k*_*off*_. We then examined whether the diffusion dynamics of EWS::FLI1 depend on RNA using fast SPT combined with the saSPT analysis method. The characteristic diffusion coefficient of freely diffusing EWS::FLI1 in the triptolide-treated cells is 1.97 ± 0.03 *μm*^2^/*s*, nearly identical to that in the DMSO-only-treated cells (Figure 4f). The bound fraction of EWS::FLI1 also does not significantly change upon triptolide treatment, with 72.48 ± 1.86% in triptolide-treated cells and 73.87 ± 2.66% in DMSO-only-treated cells, respectively (Figure 4g). These results suggest that the presence of RNA does not affect the steady-state balance of the bound and unbound EWS::FLI1 molecules in the cell nucleus. The bound fraction *f*_*bound*_is directly linked to the dissociation constant *K*_*d*_ of EWS::FLI1 binding to chromatin via

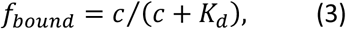

where *c* is a constant representing the intranuclear concentration of the binding sites of EWS::FLI1 on chromatin. Thus, *K*_*d*_ , the ratio of the dissociation and association rate constants (*k*_*off*_/*k*_*on*_ ), is also insensitive to the presence of RNA. Given this, and because having RNA in EWS::FLI1 hubs increases *k*_*off*_of EWS::FLI1, the presence of RNA also proportionally increases its *k*_*on*_.

Together, our results suggest that RNA plays a role in the assembly dynamics of EWS::FLI1 hubs but not their dimensions. Interestingly, nascent RNA in EWS::FLI1 hubs can increase both *k*_*on*_and *k*_*off*_of EWS::FLI1 binding to chromatin, such that their ratio is largely independent of the presence of RNA. The specific mechanism by which RNA affects the binding dynamics of EWS::FLI1 is unknown. Nascent RNA synthesized at EWS::FLI1 target genes may dynamically interact with EWS::FLI1 hubs directly through aromatic interactions^78^ and/or indirectly through RNA-binding proteins such as RNA helicase A^79,80^, allowing RNA to serve as a multivalent scaffold^81−83^ that helps concentrate EWS::FLI1 at the gene loci, thus increasing the likelihood of EWS::FLI1 binding to chromatin (increasing *k*_*on*_ ). Meanwhile, RNA might increase *k*_*off*_by destabilizing existing EWS::FLI1-chromatin interactions through charge-based repulsion or steric hindrance^33^.

### Small molecules can disrupt the nuclear distribution of endogenous EWS::FLI1

Several small molecules have been reported in the literature to affect the functions of EWS::FLI1 and some are being explored as potential therapeutic candidates. However, how these compounds affect the nuclear organization of endogenous EWS::FLI1 in Ewing sarcoma cells, especially EWS::FLI1 hub formation, remains unknown. We sought to fill this gap by examining the spatial distribution of endogenous EWS::FLI1-Halo in the knock-in A673 cells after treating the cells with these compounds using live-cell confocal imaging.

The first compound we investigated was a CDK4/6 inhibitor, LY2835219, which is reported to disrupt the condensates formed by mCherry-labeled EWS::FLI1 overexpressed in U2OS cells, partly through activating lysosomal acidification^34^. The method that identified LY2835219, DropScan, screens condensate-modulating compounds using high-content imaging of fluorescently labeled condensates in U2OS cells formed via protein overexpression. To examine the effect of LY2835219 on endogenous EWS::FLI1 hubs in Ewing sarcoma cells, we performed live-cell confocal imaging of endogenous EWS::FLI1-Halo in the knock-in A673 cells. The cells were imaged after 6 hours of treatment with 10 µM LY2835219 (stock solution in DMSO) or with DMSO only as a control (Figure 5a). While LY2835219 did not completely dissolve EWS::FLI1 hubs, it did reduce their number per cell nucleus. To quantify this difference, we calculated the percentage change in the number of puncta in the nucleus, surface roughness (standard deviation of pixel intensities across the nucleus), and area of the largest punctum in the nucleus in the cells with the LY2835219 treatment compared to the cells with the DMSO-only treatment, following an established procedure^84^ described in the Methods. We observed a statistically significant decrease in the number of puncta (Figure 5c), suggesting that LY2835219 disrupts endogenous EWS::FLI1 hubs in the native cellular context. Endogenous EWS::FLI1 hubs observed in our experiment, consistent with many previous studies^1,9,84−87^, are located in the cell nucleus, which is different from the puncta of overexpressed mCherry-EWS::FLI1 that exhibited both nuclear and cytoplasmic localizations in DropScan^34^. Regardless, our result supports the finding from DropScan that LY2835219 can disrupt EWS::FLI1 puncta. Furthermore, LY2835219 treatment was shown to significantly downregulate transcription of 20 out of 32 EWS::FLI1 target genes^34^. Wang et al. proposed that LY2835219 dissolves the condensates of overexpressed EWS::FLI1 by lysosomal activity rather than a CDK4/6 inhibition-dependent mechanism, as they ruled out the latter by showing that other selective CDK4/6 inhibitors failed to dissolve the condensates^34^. However, since many EWS::FLI1 condensates were cytoplasmic in this previous study, it remains unclear whether endogenous EWS::FLI1 hubs in the nucleus would be similarly insensitive to CDK4/6 inhibitors or be regulated by a lysosome-mediated process. Together, our findings from live-cell imaging of endogenous EWS::FLI1, combined with these previous results, highlight the therapeutic potential of LY2835219 and warrant further investigation.

**Figure 5.**
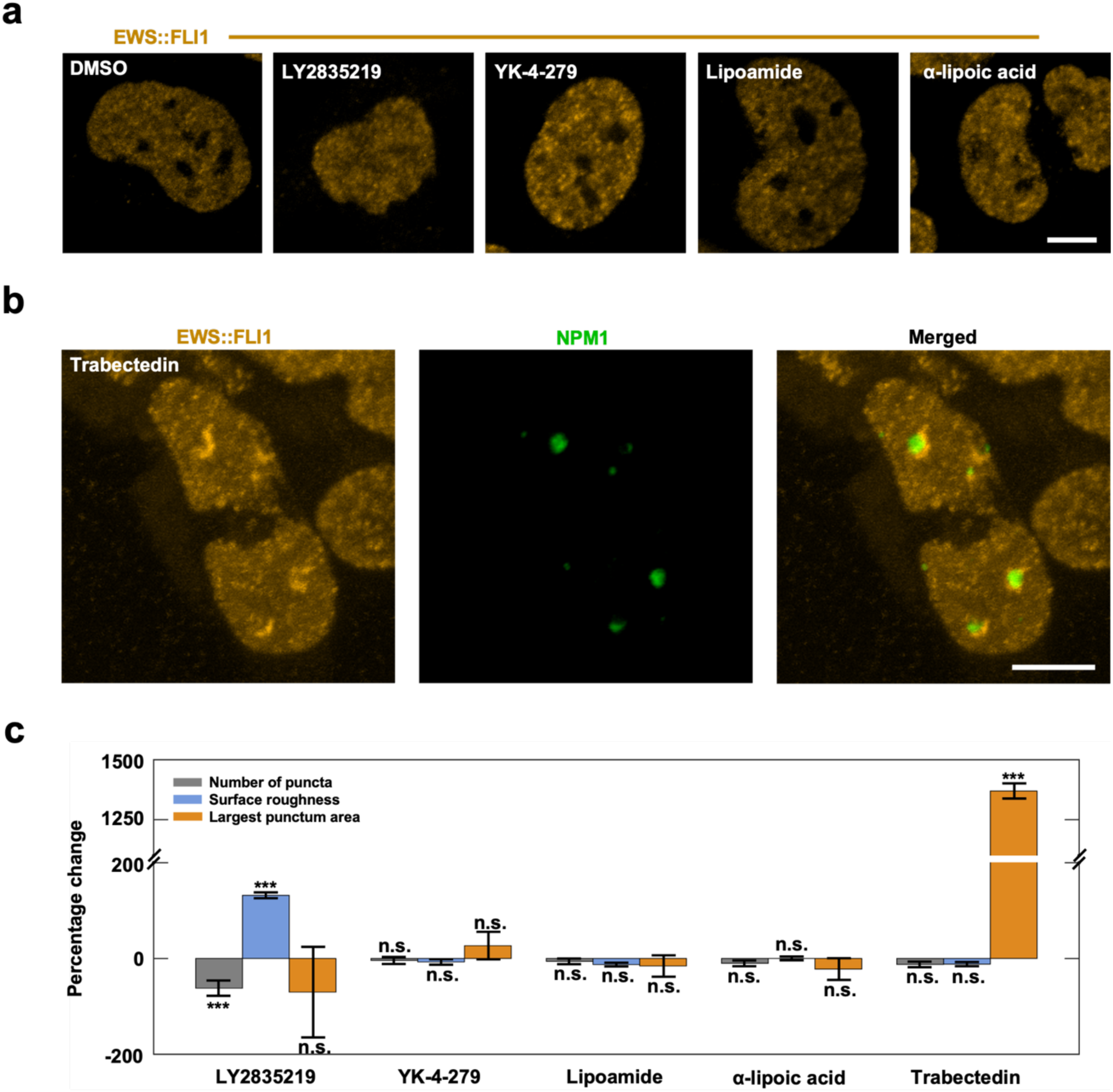
Small molecules can disrupt the nuclear organization of EWS::FLI1. (a) Live-cell confocal fluorescence images of endogenous EWS::FLI1 (labeled with the Halo ligand JFX549) in the knock-in A673 cell line after treatment with DMSO only, 10 μM LY2835219 for 6 hrs, 10 μM YK-4-279 for 15 hrs, 10 μM lipoamide for 1 hr, and 10 μM α-lipoic acid for 1 hr. (b) Live-cell confocal fluorescence images of endogenous EWS::FLI1 (labeled with the Halo ligand JFX549, yellow) in the knock-in A673 cell line transfected with mNeonGreen-labeled NPM1 that marks the nucleoli (green) and treated with 24 nM trabectedin for 1 hr followed by 23 hrs of culture in untreated media. Scale bar is 5 µm. (c) Plot of the percent change in the number of puncta, surface roughness, and largest punctum area per cell nucleus calculated as described in the Methods. Percent changes were between cells treated with each chemical and cells treated with an identical final concentration of DMSO as the solvent control. Treatment with LY2835219 reduces the number of puncta (***, *p* = 5.82 × 10^−10^, Welch’s t-test) and increases the surface roughness (***, *p* = 1.29 × 10^−6^, Welch’s t-test) in the nucleus. Treatment with trabectedin increases the size of the largest punctum area (***, *p* = 1.85 × 10^−6^, Mann-Whitney U test) in the nucleus. 18 cells were analyzed for each condition. Error bars represent SEM.

The second compound we investigated was YK-4-279, a well-characterized inhibitor of EWS::FLI1 that disrupts EWS::FLI1 interactions with RNA helicase A, leading to altered splicing and reducing transcription of EWS::FLI1 target genes, such as *UBE2C*^88^. We asked whether the effect of YK-4-279 involves disrupting the multivalent LCD-LCD interactions of EWS::FLI1 that enable its hub formation. We imaged the knock-in A673 cells expressing endogenous EWS::FLI1-Halo after treating the cells with 10 µM YK-4-279 (stock solution in DMSO) or DMSO only for 15 hours (Figure 5a). We found that the YK-4-279 treatment did not significantly change the number of puncta, surface roughness, or area of the largest punctum in the cells compared with the DMSO-only treatment (Figure 5c). This result suggests that YK-4-279 does not disrupt endogenous EWS::FLI1 hubs.

We next investigated lipoamide and α-lipoic acid, which are reported to partition into FUS droplets in vitro. The compounds also accumulate in and dissolve cytoplasmic stress granules formed upon oxidative stress, which were marked by fluorescently labeled FUS and contain all the FET family proteins (FUS, EWSR1, and TAF15)^89^. FUS is a homolog of EWSR1, and EWS::FLI1 contains the LCD of EWSR1. We thus hypothesized that lipoamide and α-lipoic acid can partition into and disrupt EWS::FLI1 hubs similarly. We imaged the knock-in A673 cells expressing EWS::FLI1-Halo after treating them with 10 µM lipoamide or α-lipoic acid (stock solutions in DMSO) or DMSO alone for 1 hour (Figure 5a). However, contrary to our hypothesis, neither lipoamide nor α-lipoic acid affected the number of puncta, surface roughness, or area of the largest punctum in the cells compared with the DMSO-only treatment (Figure 5c). The finding that neither lipoamide nor α-lipoic acid disrupts endogenous EWS::FLI1 hubs, despite the sequence similarities between the LCDs of EWS::FLI1 and FET family proteins, suggests that there are differences in the molecular mechanisms driving the formation of EWS::FLI1 hubs and cytoplasmic stress granules. This also indicates that the mechanisms by which lipoamide and α-lipoic acid mediate stress granule dissolution are independent of the LCD of EWSR1 that is present in the stress granules.

Finally, we investigated trabectedin, a DNA minor groove-binding compound that has been shown to suppress the expression of EWS::FLI1 target genes, including *NR0B1* and *CCND1*, through indirect, epigenetic mechanisms, e.g., evicting the SWI/SNF chromatin-remodeling complex from the genes, rather than through direct disruption of protein structure or interactions^90^. Previous immunofluorescence data also showed redistribution of EWS::FLI1 into the nucleolus upon trabectedin treatment, suggesting that the mislocalization of EWS::FLI1 away from its target genes may also contribute to the suppression of its transcriptional activation function. However, these experiments were performed in fixed cells^90^, and fixation can artificially change intracellular distribution of specific proteins^84^. Here, to examine the localization of endogenous EWS::FLI1 in live cells upon trabectedin treatment, we performed live-cell confocal microscopy on the knock-in A673 cells expressing EWS::FLI1-Halo after treating the cells with 24 nM trabectedin (stock solution in DMSO) or DMSO alone for 1 hour. Prior to either treatment, we incubated the cells in untreated media for 23 hours, a condition identical to the previous study^90^. However, contrary to the previous results, live-cell imaging showed that upon the trabectedin treatment, EWS::FLI1 accumulates at the periphery of the nucleolus instead of entering its interior. This localization was further confirmed by two-color imaging of endogenous EWS::FLI1-Halo and overexpressed NPM1 labeled with mNeonGreen, where the nucleolar protein NPM1 marks the location of the nucleolus (Figure 5b). Further analysis of EWS::FLI1 distribution showed that while the trabectedin treatment did not change the number of puncta and the surface roughness in the cells compared with the DMSO-only treatment, trabectedin increased the area of the largest punctum significantly (Figure 5c). According to the cell images, the largest puncta are often located on the periphery of the nucleolus. This result provides a revised view of trabectedin’s mode of action, i.e., instead of redistributing EWS::FLI1 into the nucleolus, a compartment inactive for RNA Polymerase II transcription, trabectedin enriches EWS::FLI1 at the nucleolar periphery, which is likely spatially distinct from EWS::FLI1 target genes. Our result also suggests that trabectedin does not dissolve endogenous EWS::FLI1 hubs.

Together, this set of microscopy data provides a useful resource showing how the distribution of endogenous EWS::FLI1 in live Ewing sarcoma cells can be affected by treatment with various compounds with previously reported therapeutic effects or potential. These results, together with previous publications showing functional perturbation of EWS::FLI1 by some of the examined compounds, suggest that the transcriptional activation function of EWS::FLI1 can be inhibited through diverse mechanisms, which may and may not involve disruption of its hubs.

## Discussion

An important caveat in the characterization of IDR-mediated biomolecular assemblies arises from their strong dependence on protein concentration. Numerous studies have shown that overexpressed IDR-containing proteins can form large, liquid-droplet-like puncta in the cell, which may or may not form under endogenous concentrations^21,91^. The formation of excessive condensates in such overexpression systems can distort or obscure the true properties, functions, and regulatory mechanisms of the IDR of interest. Despite this, few assembly-forming transcription regulators have been studied at strictly endogenous concentrations. Here, we fill this gap by systematically investigating the dimensions, energetics, dynamics, and regulation of endogenous EWS::FLI1 hubs in Ewing sarcoma cells using quantitative live-cell imaging methods with single-molecule resolution (Figure 6).

**Figure 6.**
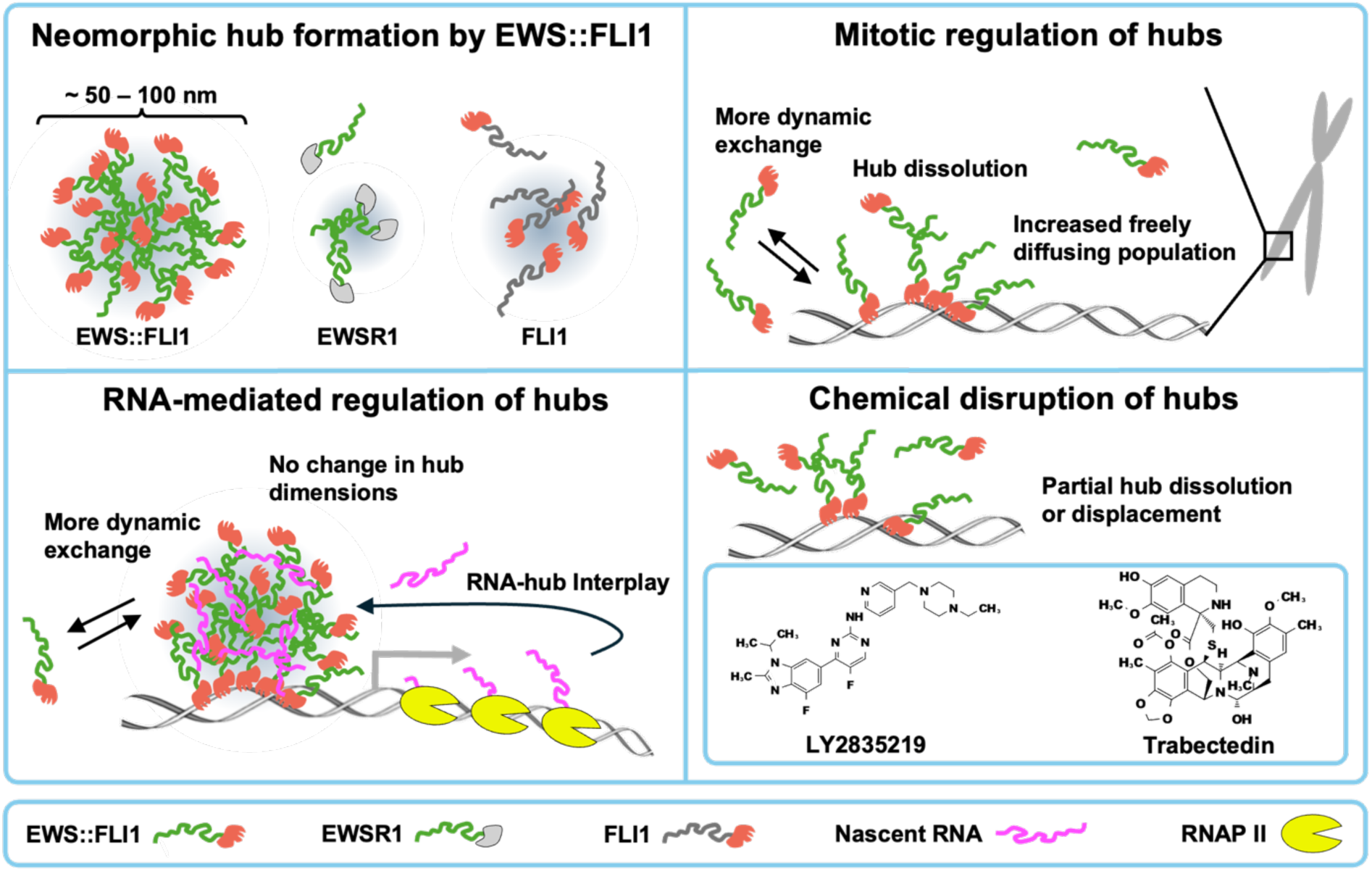
A summary of the formation, regulation, and perturbation of endogenous EWS::FLI1 hubs characterized in this work. 1) EWS::FLI1, not EWSR1 and FLI1, undergoes hub formation at native expression levels. 2) EWS::FLI1 hubs dissolve during mitosis. EWS::FLI1 retains its ability to associate with mitotic chromatin, albeit with faster binding kinetics. 3) The presence of RNA increases the in-hub binding kinetics of EWS::FLI1 without altering the hub dimensions. 4) Small-molecules LY2835219 and Trabectedin can dissolve EWS::FLI1 hubs and mislocalize EWS::FLI1 to the nuclear periphery, respectively.

Using PALM, we quantified the dimensions of sub-diffraction-limit EWS::FLI1 hubs and found that their size distribution is consistent with that of a supersaturated system predicted by the classical theory of homogeneous nucleation. While preceding studies have predicted and observed supersaturated (and subsaturated) protein clusters *in vitro*^92−97^ or in the cell^98,99^, this is the first time that the energetics and kinetics of endogenous TF hub formation have been characterized in the framework of the classical nucleation theory. Our finding raises the question of why the EWS::FLI1 hub size distribution appears to be held at a stationary nonequilibrium steady state without progressing to larger, phase-separated droplets, implying the existence of a hub dissolution mechanism. Although the identity of the dissolution pathway remains unknown, our estimated hub nucleation rate ( 34.15 hubs per second) sets the approximate timescale of the dissolution process. It will be of future interest to investigate the possible TF hub dissolution pathways that can occur at rates in this scale, such as chaperone-mediated dissolution^54,55,100^, phosphorylation-mediated dissolution^57,58,101^, and competition between multivalent interactions and DNA-protein binding^102^. In addition, our results suggest that endogenous EWS::FLI1 hubs are supersaturated clusters^8^ rather than *bona fide* LLPS droplets and are likely different from transcription condensates formed at super-enhancers reported in previous work^8,103,104^.

EWS::FLI1 exhibits multiple neomorphic behaviors that its parental proteins (EWSR1 and FLI1) lack, including remodeling GGAA microsatellites into *de novo* enhancers^105,106^, inactivating canonical ETS enhancers by displacing wild-type ETS TFs^106^, and inducing aberrant alternative splicing^107^. Here, using PALM to quantify protein distributions in the cell with sub-diffraction resolution, we showed that hub formation is another neomorphic behavior of EWS::FLI1. Specifically, we found that while both parental proteins formed puncta, neither exhibited the same degree of clustering as EWS::FLI1. This result supports our previous conclusion that hub formation behavior plays an important role in the oncogenic functions of EWS::FLI1, underscoring the potential of targeting this behavior in therapeutic development. Many oncogenic fusion proteins resulting from chromosomal translocations have been reported to undergo LLPS when overexpressed in the cell^108^. In the future, it will be interesting to investigate whether LLPS or related assembly formation is a neomorphic behavior of fusion proteins other than EWS::FLI1 and how these neomorphic assemblies may contribute to the oncogenesis of different cancers. The imaging and analysis approach we develop here will provide helpful tools for this purpose.

We then investigated whether EWS::FLI1 hubs are regulated during mitosis or by RNA using various live-cell imaging approaches. We found that during mitosis, EWS::FLI1 hubs dissolve, but EWS::FLI1 remains capable of binding to and dissociating from chromatin, albeit more dynamically and with a smaller bound fraction than during interphase. This suggest that similar to several reported TFs^75^, EWS::FLI1 can also serve as a mitotic bookmark that helps to restart transcriptional programs in daughter cells after mitosis. It will be of future interest to investigate whether and how mitotic bookmarking by EWS::FLI1 may contribute to the tumorigenesis of Ewing sarcoma. In addition, since a burst in protein phosphorylation occurs as cells go into and progress through mitosis^109,110^, our observed dissolution of EWS::FLI1 hubs during mitosis indicates a potential role of phosphorylation in regulating the hubs and warrants future investigation of this regulation mechanism.

Contrary to mitosis, when we remove nascent RNA from the cells, EWS::FLI1 binds to chromatin more stably with the hub dimensions unchanged. This finding highlights similarities and differences between EWS::FLI1 hubs and other transcription-related condensates, such as Mediator condensates, where RNA plays an essential role in condensate formation. Specifically, varying RNA concentrations strongly influences both the dimensions and dynamics of Mediator condensates, with transcriptional inhibition increasing both condensate size and lifetime. In contrast, EWS::FLI1 hubs are predominantly maintained by protein-DNA and protein-protein interactions, with RNA contributing only to regulating EWS::FLI1 binding kinetics. The differential roles of RNA in the regulation of Mediator condensates and EWS::FLI1 hubs could be due to differences in the chemistry underlying the molecular interactions that drive the formation of the respective assemblies. The IDR of the Mediator subunit MED1 contains alternating blocks of charged amino acids that drive condensate formation^3,111^. RNA, a highly-charged polyanion, also interacts with the MED1 IDR primarily via electrostatic interactions, thus strongly coupling the formation and dissolution of MED1 condensates to local RNA concentration^33^. On the other hand, EWS::FLI1 hub formation is driven mainly by interactions between aromatic residues, particularly tyrosines in the EWS LCD^1^. These interactions are less sensitive to changes in bulk electrostatics ^33,112^, thus resulting in a lower impact of local RNA concentration on EWS::FLI1 hub assembly. Our observations are also related to previous work showing that an increase in the RNA-protein ratio can reduce the ability of EWSR1 and FUS, members of the FET family of RNA-binding proteins^113^, to form condensates. Notably, this effect was previously observed when the proteins were artificially driven to phase separation via overexpression. In contrast, our measurements of EWS::FLI1 under endogenous expression levels suggest that RNA regulates only the binding dynamics of EWS::FLI1, without altering its hub size.

Finally, we showed that in live Ewing sarcoma cells, LY2835219 partially dissolves endogenous EWS::FLI1 hubs and trabectedin redistributes EWS::FLI1 to the periphery of the nucleolus, away from its target genes. This is a useful finding given that recent work indicates that assemblies of disease-causing TFs are potential alternative therapeutic targets, owing to the notoriously “undruggable” nature of most TFs, which is due to their largely disordered nature and lack of well-defined binding pockets. For EWS::FLI1 specifically, our earlier work shows that its hub formation mediated by multivalent LCD-LCD interactions plays an essential role in oncogenic transcription in Ewing sarcoma ^1,9^, suggesting disruption or mislocalization EWS::FLI1 hubs as a potential therapeutic strategy. Here, our finding that LY2835219 and trabectedin can alter the nuclear distribution of endogenous EWS::FLI1 further supports the previously reported therapeutic potential of the compounds.

Overall, our study combined CRISPR/Cas9-mediated genome editing and quantitative live-cell and single-molecule imaging techniques to characterize the formation and regulation of endogenous TF hubs at an unprecedented resolution. We quantified the dimensions of EWS::FLI1 hubs along with the energetics and kinetics of hub nucleation. We uncovered the neomorphic nature of EWS::FLI1 hubs and dynamic hub regulation during mitosis and by RNA. We also showed that endogenous EWS::FLI1 hubs in Ewing sarcoma cells can be disrupted by small molecules, including LY2835219 and trabectedin, suggesting a route for therapeutic intervention. While our study focuses on a single TF EWS::FLI1, the findings we made here, particularly those related to the formation and regulation of TF hubs, may apply to diverse TFs, including regular and pathological ones. Future studies applying similar approaches to a broad range of TFs will inform the generalizability of our findings.

## Methods

### Cell Culture

The knock-in A673 cell line expressing endogenous EWS::FLI1-Halo from Chong et al.^1^ was grown in 4.5 g/L high-glucose DMEM (Thermo Fisher, 10566016) with 10% FBS (Fisher Scientific, SH3039603) and 1% penicillin-streptomycin (Thermo Fisher, 15140122). For live-cell imaging, the phenol-red-free version of high-glucose DMEM (Thermo Fisher, 31053028) was used instead. The human U2OS osteosarcoma cell line was cultured in 1 g/L low-glucose DMEM (Thermo Fisher, 10567014) with 10% FBS and 1% penicillin-streptomycin. Cells were synchronized using nocodazole (Sigma Aldrich, M1404-2MG). All cell lines were cultured in a humidified incubator at 37°C and 5% CO_2_.

### Single-molecule imaging

All single-molecule imaging was performed on 25 mm #1.5H coverslips (Marienfeld Superior, 0117650). Coverslips were cleaned by immersing in 1 M KOH and sonicating for 1 hr, rinsing thrice with double-distilled water, immersing in pure ethanol and sonicating for 1 hr, then immersing in 20% ethanol diluted in double-distilled water for storage.

Single-molecule imaging was performed on a Nikon Eclipse Ti2 N-STORM microscope setup with a x100/NA 1.49 oil-immersion objective (Nikon, CFI SR HP Apochromat TIRF 100XAC Oil, MRD01995). Samples were illuminated under a highly inclined and laminated optical sheet (HILO)^114^ with a Nikon LUN-F solid-state laser unit using the following lasers: 405 nm (photoactivation of PA-JF549 and PA-JF646^115^ and excitation of Hoechst 33342), 561 nm (excitation of JFX549 and PA-JF549), and 640 nm (excitation of JFX646^116^ and PA-JF646). For single-channel experiments, fluorescence from JFX549 and PA-JF549 was filtered through a 593/40 nm single-band bandpass filter (Semrock, FF01-593/40-25) and fluorescence from JFX646 and PA-JF646 was filtered through a 676/37 nm single-band bandpass filter (Semrock, FF01-676/37-25) and detected on an iXon Ultra 897 EMCCD (Andor). For two-channel experiments, the emission fluorescence was first split using a 635 nm single-edge dichroic beamsplitter (Semrock, Di02-R635-25×36) before being similarly filtered through the appropriate single-band bandpass filters and detected on iXon Ultra 897 EMCCDs. For live-cell imaging, a stage incubator was used to maintain humidity and 37°C with 5% CO_2_. Fixed-cell imaging was performed at room temperature. All hardware was controlled through the NIS-Elements software (Nikon).

### Photoactivated localization microscopy (PALM)

Cells were cultured on cleaned coverslips then stained with 200 nM PA-JF549 following the protocol described in Yoshida et al.^117^. Cells were then fixed using 4% paraformaldehyde (VWR, 100503-917) supplemented with 0.2% glutaraldehyde (Sigma Aldrich, 340855-25ML) for 15 minutes, then rinsed thrice with 1X PBS for 5 min (Thermo Fisher, 18912014). Coverslips were then transferred into Attofluor Cell Chambers (Thermo Fisher, A7816). 0.1 μm yellow-green Invitrogen Fluospheres (Thermo Fisher, F8803) were diluted in DPBS with calcium and magnesium (Thermo Fisher, 14-040-141) and added to the coverslips for 10 minutes at room temperature to allow the beads to adsorb to the coverslip.

The samples were imaged on the single-molecule microscope described above under continuous, simultaneous photoactivation (405 nm) and excitation (561 nm). The 405 nm laser intensity was gradually increased over the course of the acquisition to maintain an approximately constant number of photoactivated molecules, and the 561 nm laser was held constant. A 50 ms integration time was used and cells were imaged until all single molecules were bleached (typically at least 24000 frames). Images of the fiducial marker positions were obtained every 100 frames.

NIS-Elements was used to identify single-molecule localizations in each frame, with the following settings: minimum height: 1500, maximum height: 65535, CCD baseline: 100, minimum width: 200 nm, maximum width: 600 nm, initial fit width: 300 nm, maximum axial ratio: 1.2, maximum displacement: 0. Fiducial markers were identified under identical settings except for minimum height: 3000. Drift correction was performed based on the fiducial marker positions, and the localizations were exported as a .csv file.

Even when using well-behaved fluorophores, single-molecules typically appear in multiple frames due to a combination of variable photobleaching times and photoblinking^118^. More specifically, rather than photobleaching permanently after a fixed amount of time, fluorophores can enter a temporary dark state from which they recover and reappear, often “photoblinking” a number of times before permanently bleaching. This is especially problematic in the characterization of TF hubs because it becomes difficult to distinguish between the contributions of true TF clustering and blinking. Thus, to minimize these artifacts, endogenous EWS::FLI1-Halo was labeled with the photoactivatable ligand PA-JF549 which features minimal blinking^119^, and the PALM acquisitions were performed with low intensity photoactivation to minimize the probability of multiple molecules within a diffraction-limited spot fluorescing simultaneously. Even with these strategies, it is still necessary to correct for multiple appearances of the same protein, motivating pair-correlation PALM (PC-PALM) and our photoblinking correction.

Raw localizations were analyzed using a custom python implementation of PC-PALM^40,46,120^. While PC-PALM provides an excellent way of discriminating between localization clusters due to true protein clusters and due to photoblinking (particularly for emitters with high blinking rates or long temporary dark states), it cannot identify which localizations correspond to a single protein, i.e. it cannot recover the set of true protein localizations. Further, it is particularly sensitive to edge effects and variable localization densities, thus only a subset of cellular localizations (corresponding to a representative, relatively homogeneous subregion of the nucleus) can be analyzed. In order to generate a reconstructed image with true localizations, we had to directly filter out the localizations due to photoblinking.

Often, the time between reappearances of a single emitter is much shorter than the amount of time before another emitter is excited in the same position within what is expected with localization error and previous work has shown that identifying spatiotemporal clusters can be an effective way to identify photoblinking artifacts^121^. Existing strategies often choose a threshold cutoff time for either the time until photobleaching, *t*_*bleach*_, or the time between appearances, *t*_*off*_. For the *t*_*bleach*_ strategy, all localizations within the threshold time of an initial localization are assumed to have come from the same molecule, and their positions are averaged and they are treated as one molecule. For the *t*_*off*_strategy, after localization disappears, a reappearance that takes place within the threshold time elapsing is assumed to have come from the same molecule, and the localization are similarly averaged. The specific threshold values used, whether to use the *t*_*off*_ or *t*_*bleach*_ strategy, and whether either of these strategies is suitable at all, are all dependent on the fluorophore used. Thus, we first needed to characterize the blinking behavior of PA-JF549.

Many groups have investigated the blinking properties of fluorophores by purifying fluorescent proteins and tethering them to coverslips^40^. However, the blinking properties of fluorophores is also well known to be highly environment dependent^122^. Thus, we chose to examine the blinking of PA-JF549 in fixed, A673 cells stably expressing Halo-NLS, which does not significantly cluster. Other than staining concentration, the samples were prepared and imaged under identical conditions as our PALM experiments. The cells were stained with 0.5 nM of PA-JF549, making it highly unlikely that any Halo-NLS bound by a ligand would appear in the same location as another ligand-bound molecule. This ensured that any localizations in a particular region would all be due to a single PA-JF549 molecule. We then performed PALM imaging across ten cells, and in each cells the localizations were clustered using DBSCAN (ε = 10 nm, minimum points = 1). We confirmed that PA-JF549 features extremely low blinking, appearing on average 1.12 times, on par with an existing estimate from the literature^119^. Next, we adopted a strategy similar to those of Annibale et al.^121^ and Lee et al.^123^ for determining the appropriate *t*_*off*_ threshold value (hereafter, 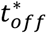) using our Halo-NLS data. Given the total number of localizations within a region *n*_*total*_, the true number of proteins *n*_*true*_, and the mean number of blinks 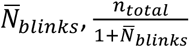 becomes an excellent estimator of *n*_*true*_for increasing *n*_*total*_as long as 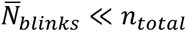. Thus, we can determine an appropriate value of 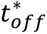 by estimating *n*_*true*_ across all Halo-NLS cells using *n*_*total*_ and the observed value of 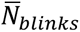 then filtering blinking using different values of 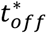 and totaling the number of “true” localizations, 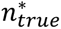. until 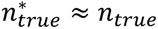. We did this and found that *t*_*off*_ = 0.05 ms, that is tolerating a single empty frame between blinking events, was sufficient to recover 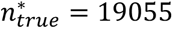 of *n*_*true*_ = 19079 total Halo-NLS localizations, corresponding to a relative error of 0.13%. We then applied this threshold value to our raw PALM localizations.

To visualize whether there was a significant difference in the spatial distribution of proteins in different PALM datasets, we computed Ripley *L*(*r*) − *r* for each cell over values of *r* from 0 to 750 nm in Python using the ‘RipleysKEstimator’ function from the ‘astropy.stats’ package^124^. The area of each nucleus was estimated by generating an alpha shape around the localizations ( α = 2 ) using the ‘alphashapè package. While spatial descriptive statistics provides an excellent visualization of which proteins are more clustered than others and over what length scales, a great deal of information on the single cluster level is lost. Thus, we identified individual hubs by clustering the raw localizations using DBSCAN (ε = 10 nm, minimum points = 3), confirming by eye the suitability of these parameters. For each hub, the diameter was calculated as the maximum pair-wise distance between localizations and the “mode” diameter was extracted by approximating the hub diameter distribution as lognormal.

Our analysis investigating the hub distribution through the lens of homogeneous nucleation was based on the framework established by Naranayan et al.^125^. Measurements of both critical radius *R*_*c*_and free energy barrier to nucleation Δ*G*_*c*_are insensitive to constant multiplicative factors in hub “size” *n*^125^, thus 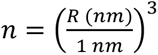 was calculated for each hub with diameter, *d*, larger than our localization precision, ∼20 nm. Here, *n* is proportional to the number of EWS::FLI1 molecules in each hub. The offset factor is primarily due to the fact that every EWS::FLI1-Halo molecule does not end up being detected; literature measurements of the effective labeling efficiency of PA-JF549 place it at around 21%^119^. Eqs. (1) and (2) are valid only for subcritical hubs, thus only hubs with *n* ≤ 1 × 10^6^ were analyzed. This condition ensured that only subcritical hubs, accounting for over 85% of hubs in each cell, were included. Then *P*(*n*) was generated using bins of size ∼5000 and then − log *P*(*n*) was calculated and fit as − log *P*(*n*) = *ɛ*^′^*n*^2/3^ − Δ*μ*^′^*n* − *b*. Fit values of *ɛ*^′^ and Δ*μ*^′^ were used to calculate *R*_*c*_ and Δ*G*_*c*_ as described in Supplementary Note A. Estimation of the nucleation rate, *J*, is described in Supplementary Note B.

### Slow single-particle tracking (SPT)

Slow SPT was performed following Yoshida et al.^71^. Briefly, cells were cultured on cleaned coverslips then for slow SPT during mitosis, cells were stained with 20 nM PA-JF646 and DNA was labeled with Hoechst 33342. For slow SPT after transcriptional inhibition, cells were stained with 20 nM PA-JF646 and 100 nM JFX549. This enabled the simultaneous visualization of individual EWS::FLI1 molecules in the PA-JF646 channel against the real-time distribution of DNA or EWS::FLI1 hubs. Coverslips were then transferred into Attofluor Cell Chambers and DMEM was exchanged for phenol-red-free DMEM.

The samples were then imaged on the single-molecule microscope described above. Single molecules were excited using the 640 nm laser during the exposure time and they were photoactivated using the 405 nm laser during the transition time between exposures. 500 ms exposure times were used such that mobile particles were blurred out. The 405 nm laser intensity was gradually increased over the course of the acquisition to maintain an approximately constant number of photoactivated molecules, and the 640 nm laser was held constant. For simultaneous single-molecule imaging with DNA, one frame of the Hoechst 33342 channel was taken before and after acquisition. For simultaneous single-molecule imaging with EWS::FLI1 hubs, every 10 seconds, one frame of the bulk EWS::FLI1 channel was taken. One the timescale of the acquisition, DNA is not that mobile, thus before and after images are sufficient for a timelapse. EWS::FLI1 hubs remain mobile, thus the need for more frequent timelapse images. For each movie, 2000 frames were acquired, and movies were split into single-molecule channels (PA-JF646) and bulk channels (Hoechst 33342 or JFX549).

Single-molecules were localized and assembled into trajectories using SLIMfast^126^, a MATLAB-based GUI implementation of the MTT algorithm^127^. The following parameters were used: localization error: 10^−6^, deflation loops: 3, maximum number of competitors: 5, maximum expected diffusion coefficient: 0.1 µm^2^/*s* . Using the bulk channels, real-time binary masks of DNA/EWS::FLI1 hub position were generated with custom ImageJ macros from Chong et al.^1^. By changing the threshold for EWS::FLI1, masks of both EWS::FLI1 hubs and the whole nucleus were created. Then, using different custom ImageJ macros also from [], trajectories were sorted into those overlapping with DNA/EWS::FLI1 hub masks and those not. The trajectories overlapping with masks represent stable binding to DNA/EWS::FLI1 hubs. Survival curves defined as 1 − *CDF*, i.e. the complement of the cumulative distribution function, were generated from the overlapping trajectories and fit to the following two-component exponential model.

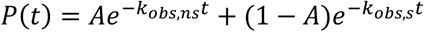

Here, A is the fraction of non-specifically bound molecules, *k*_*obs*,*ns*_is the observed dissociation rate of nonspecifically bound molecules, and *k*_*obs*,*s*_is the observed dissociation rate of specifically bound molecules. The observed dissociate rate of specifically bound molecules is the sum of the photobleaching rate, *k*_*pb*_, and the true specific dissociation rate, *k*_*true*,*s*_, since both dissociation and photobleaching can result in a trajectory ending. The photobleaching rate is obtained by doing slow SPT under identical settings on an A673 cell line stably expressing H2B-Halo described in Chong et al.^1^ and extracting the *k*_*obs*,*s*_for H2B, which rarely dissociates on the timescales measured by slow SPT. The mean residence time, *τ̅*_*s*_, is then given as

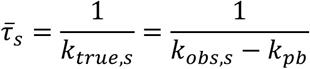

### Fast single-particle tracking (SPT)

Cells were cultured on cleaned coverslips then stained with 20 nM PA-JF646. Coverslips were then transferred into Attofluor Cell Chambers and DMEM was exchanged for phenol-red-free DMEM.

The samples were then imaged on the single-molecule microscope described above under stroboscopic illumination based on Chong et al.^9^. Cameras were synchronized at the beginning of each acquisition with 0.3 µs shift speeds and +4 vertical clock voltage amplitudes enabling 5 ms exposure times with 158.01 µs transition times. Single molecules were excited using the 640 nm laser during only the first 1 ms of each exposure. Single molecules were photoactivated using the 405 nm laser during the transition times. The 405 nm laser intensity was gradually increased over the course of the acquisition to maintain an approximately constant number of photoactivated molecules, and the 640 nm laser was held constant. For each movie, 20000 frames were acquired.

Single molecules were localized and assembled into trajectories using SLIMfast^126^, using the following parameters: localization error: 10^−5^, deflation loops: 2, maximum number of competitors: 3, maximum expected diffusion coefficient: 3.5 µm^2^/*s*. The resulting trajectories were then analyzed using the state array SPT (saSPT) python package developed in Heckert et al.^128^. The state array was defined over one hundred logarithmically spaced diffusion coefficients from 10^−2^ to 10^1^ µm^2^/*s*, inclusive, and twenty linearly spaced localization errors from 0 to 0.07 µm, inclusive. The posterior occupations were then marginalized over localization error to obtain the diffusion coefficient distribution. Bound fractions were computed as the sum of the occupation values of diffusion coefficients under 0.1 µm^2^/*s* corresponding to stably bound molecules.

### Live-cell confocal microscopy

Cells were cultured on 35 mm MatTek glass-bottom dishes (MatTek, P35G-1.5-20-C) and stained with 200 nM JFX549 or JFX646. For two-color imaging during mitosis, DNA was labeled with Hoechst 33342. For two-color imaging with the nucleolus, cells were transfected with the mNeonGreen-NPM1 construct 24 hrs before imaging using Lipofectamine 3000 (Fisher Scientific, L3000001) following the manufacturer’s protocol. DMEM was then replaced with phenol-red-free DMEM.

Confocal imaging was performed on a Nikon Eclipse Ti2 AX R point-scanning confocal microscope setup with a x60/NA 1.42 (Nikon, CFI Plan Apochromat Lambda D 60X Oil, MRD71670). Samples were illuminated with a Nikon LUA-S4 laser unit using the following lasers: 405 nm (excitation of Hoechst 33342, (VWR, PI62249)), 488 nm (excitation of mNeonGreen), 561 nm (excitation of JFX549^116^ and Quasar 570), and 640 nm (excitation of JFX646^116^ and Quasar 670). For multichannel images, emission filters and detection sequence were chosen such that there was no bleed-through between the channels. Z-stacks were acquired with slice intervals of 0.5 µm and a pinhole size of 1 Airy unit. A stage incubator was used to maintain humidity and 37°C with 5% CO_2_.

### RNA fluorescence in situ hybridization (RNA FISH)

Knock-in A673 cells were plated on 18 mm circular No 1.5 coverslips (Electron Microscopy Sciences, 72222-01) cleaned by soaking with 70% ethanol. Cells were then stained with 200 nM JFX549 or JFX646 chosen such that the labeled EWS::FLI1-Halo is spectrally distinct from the RNA FISH probe. Intron RNA FISH was performed following the protocol for adherent cells published online by Stellaris using FISH probes targeting the introns of *ABHD6* and *CAV1* from Chong et al.^9^ designed using the online Stellaris Probe Designer software. The *ABHD6* probe was conjugated with Quasar 670 and *CAV1* probe was conjugated with Quasar 570. Samples were then imaged at room temperature on the confocal system described above.

### CRISPR/Cas9-mediated genome editing

We created the knock-in A673 cell line expressing endogenous Halo-EWSR1 using published procedures^129^, except that we exploited the HaloTag for fluorescence-activated cell sorting (FACS). We first co-transfected A673 using a repair plasmid together with the Cas9 plasmid (3:1 mass ratio for repair plasmid to Cas9 plasmid). For the repair vector, we modified a pUC57 plasmid to incorporate coding sequences for TEV protease recognition epitope (EDLYFQS), HaloTag and FLAG-tag flanked by ∼750 bp of genomic homology sequence (homology arm) of EWSR1 N-terminus on either side. The TEV linker sequence was between HaloTag and EWSR1. We introduced synonymous mutations in the homology sequence, where necessary, to prevent the Cas9-sgRNA complex from cutting the repair vector.

We designed two sgRNAs (CGGGTGAGTATGGTGGAACT and GGAGAGAAAATGGCGTCCAC) using the Zhang lab CRISPR design tool (http://tools.genome-engineering.org), cloned them into the Cas9 plasmid, and transfected A673 cells with the Cas9-sgRNA plasmid and the repair vector. 24 hours after transfection, we pooled cells transfected with each sgRNA individually and FACS-sorted for YFP (mVenus)-positive, successfully transfected cells. To generate the HaloTag knock-in, we grew the YFP-positive cells for 19 days, labeled them with 500 nM HaloTag TMR ligand (Promega, G8251), FACS-sorted for HaloTag-positive cells, and plated one cell per well into 96-well plates. Clones were then expanded and screened by genomic PCR using a genomic primer external to the homology sequence and an internal HaloTag primer. Successfully edited clones were further verified with western blot. EWSR1 shares the same N-terminus with EWS::FLI1. We chose a clone with HaloTag knock-in at EWSR1 and normal expression of EWS::FLI1 for further studies.

### Stable cell line construction

The A673 cell line stably expressing Halo-NLS was generated using PiggyBac transposition and drug selection. Briefly, the cDNA encoding Halo-NLS was cloned into a PiggyBac vector co-expressing a G418 resistance gene, and this vector was co-transfected together with a SuperPiggyBac transposase vector into the A673 cell line using Fugene 6 (Promega, E2691) according to the manufacturer’s instructions. 48 hours after transfection, we started with selection by adding 0.5 mg/ml G418. Untransfected A673 cells were treated with G418 in parallel, and selection was judged to be complete once no untransfected cells remained (∼ 1 week). The cells were further selected for 2 weeks under 0.25 mg/ml G418 before stocks were frozen.

### Western blot

Cells were washed with ice-cold PBS and then lysed in mild lysis buffer containing protease inhibitor cocktail (Millipore Sigma, 11836170001), 50 mM Tris (pH = 7.5), 150 mM NaCl, 0.5 % Triton X-100. Total cell lysates were incubated on ice for 30 min and then centrifuged at 15,000g at 4°C for 20 min. Equal amounts of total protein were separated via 4-15% SDS-PAGE and transferred to nitrocellulose membranes. The membranes were blocked with TBS containing 0.1% Tween-20 and 5% milk and then incubated with primary antibodies overnight at 4°C. The next day, membranes were washed, incubated with the appropriate horseradish peroxidase-conjugated secondary antibodies, and were developed using horseradish peroxidase-based chemiluminescent substrate (Thermo Scientific, 34580). The following primary antibodies were used for immunoblotting: EWSR1 antibody (Abcam, ab133288), GAPDH antibody (Cell Signaling Technology, 2118S), horseradish peroxidase-labeled goat anti-rabbit secondary antibodies (Invitrogen, A21245).

### Statistical analyses

For normally distributed data with equal variance, Student’s t-tests were used. For normally distributed data with unequal variance, Welch’s t-tests were used. For skew data, Mann-Whitney U test was used. All statistical tests were performed in Python using the ‘scipy.stats’ package. Significance is abbreviated in figures according to the following key: n.s. is not statistically significant, * is p < 0.05, ** is p < 0.01, and *** is p < 0.001.

## Acknowledgements

We thank Dr. Luke Lavis for providing the fluorescent HaloTag ligands. This work was supported by the National Science Foundation Graduate Research Fellowship Program under grant number DGE-1745301 (S.Y.), the National Institutes of Health under grant number GM043974 (W.G.D), the Pew-Stewart Scholar for Cancer Research Award (S.C.), the Searle Scholar Award (S.C.), the Merkin Innovation Seed Grant (S.C.), the Mallinckrodt Research Grant (S.C.), the Margaret E. Early Medical Research Trust 2024 Grant (S.C.), and the Alex’s Lemonade Stand Foundation Innovation Grant under award number 1260879 (S.C.).

## Author Contributions

S.C. and S.Y. conceived the study, designed the experiments, and interpreted the results. S.C. constructed all knock-in cell lines. S.Y. performed all imaging experiments, analysis, and physical modeling. Y.Z. performed western blots. W.D. and A.K. provided reagents and advised on cell cycle experimental design and interpretation. S.Y. prepared the original draft and S.C. and S.Y. reviewed and edited the manuscript. S.C. supervised the work and acquired funding.

## Materials Availability Statement

The materials described in this study are available on request.

