## Supporting Information for "Dynamic regulation of endogenous transcription factor hubs at single-molecule resolution"

Shawn Yoshida[1,2], Yanghao Zhong[1], Akiko Kumagai[2], William G. Dunphy[2], Shasha Chong[1]*

[1] *Division of Chemistry and Chemical Engineering, California Institute of Technology, Pasadena, CA 91125, USA*

[2] *Division of Biology and Biological Engineering, California Institute of Technology, Pasadena, CA 91125, USA*

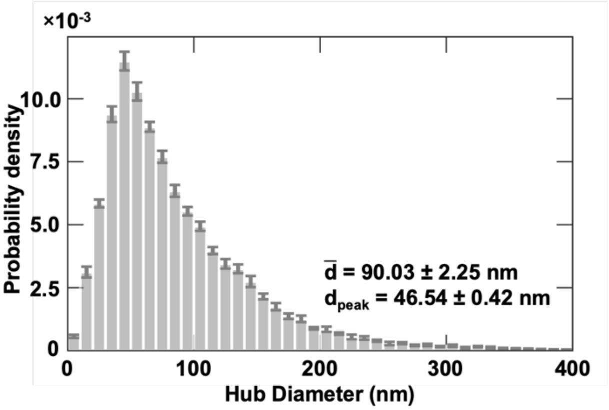


**Figure S1. PALM microscopy characterizes the dimensions of EWS::FLI1 hubs after treatment with triptolide.** The distribution of mean hub diameters generated by clustering photoblinking-corrected localizations with DBSCAN.

**Supplementary Note A:**

In the mean-field treatment of homogeneous nucleation, the free energy corresponding to the nucleation of a spherical hub of radius, $R$, is written as

$$\Delta G = 4\pi R^2\gamma - \frac{4}{3}\pi R^3 \nu^{-1} k_B T \log(S) \qquad \text{(S1)}$$

where $\gamma$ is the surface energy at the interface, $\nu$ is the volume occupied by a single molecule, and $S = c/c^*$ is the supersaturation, or the ratio of the actual concentration to the equilibrium or saturation concentration. Then, for $n$ molecules, the total volume occupied by the hub is $n\nu = \frac{4}{3}\pi R^3$ and rearranging to obtain expression for $R$ and substituting it into Equation S1 yields

$$\Delta G = (36\pi)^{1/3}\nu^{2/3}\gamma n^{2/3} - nk_B T \log(S) \qquad \text{(S2)}$$

Defining $\varepsilon \coloneqq (36\pi)^{1/3}\nu^{2/3}\gamma$ and recognizing that $\Delta\mu = k_B T \log(S)$, we obtain Equation 1, the compact form of Equation S1 used in the main text, Equation 1.

$$\Delta G = \varepsilon n^{2/3} - \Delta\mu n \qquad \text{(S3)}$$

The critical hub size $n_c$, the corresponding critical radius $R_c$, and free energy barrier to nucleation $\Delta G_c$, are set by the maximum of the free energy profile; thus, we set $\frac{d}{dn}\Delta G = 0$ and find $n_c = \left(\frac{2\varepsilon}{3\Delta\mu}\right)^3$, corresponding to

$$R_c = \frac{2\varepsilon}{3\Delta\mu} \quad \text{(S4)}$$

and

$$\Delta G_c = \frac{4\varepsilon^3}{27\Delta\mu^2} \quad \text{(S5)}$$

**Supplementary Note B:**

Assuming Becker and Döring kinetics where individual molecules (rather than multimers) join and leave hubs, the steady-state nucleation rate per unit volume follows the Zeldovich form given by[1]

$$J = \rho\beta Z exp\left(\frac{-\Delta G_c}{k_B T}\right) \quad \text{(S6)}$$

Where $\rho$ is the number density of molecules in $m^{-3}$, $\beta$ is the impingement rate of molecules in $s^{-1}$, and $Z$ is the Zeldovich factor, which accounts for the curvature of the free energy profile around the critical radius.

We can estimate that $\rho = 3.20 \times 10^{19}\ m^{-3}$ using 200 nM the concentration of EWS::FLI1 from Chong et al. 2018[2] with Avogadro's number. The impingement rate for diffusion-limited (rather than reaction-limited) incorporation can be estimated as $\beta = 4\pi R_c D\rho = 77.98\ s^{-1}$, where $D = 1.97 \times 10^{-12}\ m^2 s^{-1}$ is the diffusion coefficient of EWS::FLI1 from the main text. The Zeldovich factor is a function of the curvature of $\Delta G$ at the critical size and is given as

$$Z = \sqrt{\frac{-1}{2\pi k_B T}\left(\left.\frac{\partial^2 \Delta G}{\partial n^2}\right|_{n=n_c}\right)} \quad \text{(S7)}$$

We can rewrite this using Equation 1 yielding

$$Z = \frac{1}{3k_B T}\pi^{-1/2}\varepsilon^{1/2}n_c^{-2/3} \quad \text{(S8)}$$

Notably, $Z$ (hence $J$) is not insensitive to constant multiplicative factors of $n_c$. Thus, rather than using the fit value, we take the average number of localizations in hubs of size $R_c$ and divide by 0.21, the effective localization efficiency for PA-JF549 to estimate that $n_c = 153.21$. Using this value of $n_c$ together with the fitted value $\varepsilon = 0.0015 k_B T$ yields $Z = 2.54 \times 10^{-4}$. We can also use the fitted value of $\frac{\Delta G_c}{k_B T} = 4.88$. Using our measurements with Equation S6, we estimate that $\mathrm{J} = 6.83 \times 10^{16}\ m^{-3} s^{-1}$.
